# *Tex19.1* Regulates Meiotic DNA Double Strand Break Frequency in Mouse Spermatocytes

**DOI:** 10.1101/099119

**Authors:** James H. Crichton, Christopher J. Playfoot, Marie MacLennan, David Read, Howard J. Cooke, Ian R. Adams

## Abstract

Meiosis relies on the SPO11 endonuclease to generate the recombinogenic DNA double strand breaks (DSBs) required for homologous chromosome synapsis and segregation. The number of meiotic DSBs needs to be sufficient to allow chromosomes to search for and find their homologs, but not excessive to the point of causing genome instability. Here we report that meiotic DSB frequency in mouse spermatocytes is regulated by the mammal-specific gene *Tex19.1*. We show that the chromosome asynapsis previously reported in *Tex19.1^-/-^* spermatocytes is preceded by reduced numbers of recombination foci in leptotene and zygotene. *Tex19.1* is required for the generation of normal levels of *Spo11*-dependent DNA damage during leptotene, but not for upstream events such as MEI4 foci formation or accumulation of H3K4me3 at recombination hotspots. Furthermore, we show that mice carrying mutations in the E3 ubiquitin ligase UBR2, a TEX19.1-interacting partner, phenocopy the *Tex19.1^-/-^* recombination defects. These data show that *Tex19.1* and *Ubr2* are required for mouse spermatocytes to generate sufficient meiotic DSBs to ensure that homology search is consistently successful, and reveal a hitherto unknown genetic pathway regulating meiotic DSB frequency in mammals.

**Author Summary:** Meiosis is a specialised type of cell division that occurs during sperm and egg development to reduce chromosome number prior to fertilisation. Recombination is a key step in meiosis as it facilitates the pairing of homologous chromosomes prior to their reductional division, and generates new combinations of genetic alleles for transmission in the next generation. Regulating the amount of recombination is key for successful meiosis: too much will likely cause mutations, chromosomal re-arrangements and genetic instability, whereas too little causes defects in homologous chromosome pairing prior to the meiotic divisions. This study identifies a genetic pathway requiredto generate robust meiotic recombination in mouse spermatocytes. We show that male mice with mutations in *Tex19.1* or *Ubr2*, which encodes an E3 ubiquitin ligase that interacts with TEX19.1, have defects in generating normal levels of meiotic recombination. We show that the defects in these mutants impact on the recombination process at the stage when programmed DNA double strand breaks are being made. This defect likely contributes to the chromosome synapsis and meiotic progression phenotypes previously described in these mutant mice. This study has implications for our understanding of how this fundamental aspect of genetics and inheritance is controlled.

## Introduction

Recombination plays key roles in meiosis and gametogenesis through facilitating the pairing and reductional segregation of homologous chromosomes, and by increasing genetic variation in the next generation. Meiotic recombination is initiated when programmed DNA double strand breaks (DSBs) are generated during the leptotene stage of the first meiotic prophase. Meiotic DSBs recruit a series of recombination proteins visualised cytologically as recombination foci, and initiate a search for homologous chromosomes thereby promoting homologous chromosome synapsis during zygotene. Recombination foci continue to mature while the chromosomes are fully synapsed in pachytene, and eventually resolve into crossover or non-crossover events. Crossovers exchange large tracts of genetic information between parental chromosomes, increasing genetic diversity in the population. Furthermore, these crossovers, which physically manifest as chiasmata, hold homologs together after they desynapse in diplotene and help to ensure that homologous chromosomes undergo an ordered reductional segregation at anaphase I [1,2].

Meiotic DSBs have a non-random distribution across the genome, and their frequency and locationplay an important role in shaping the recombination landscape [2,3]. In male mice, a few hundred meiotic DSBs are generated during leptotene, around 20-25 of which mature into crossovers. The positions of meiotic DSBs across the genome are determined by PRDM9, a histone methyltransferase that mediates trimethylation of histone H3 lysine 4 (H3K4me3) at recombination hotspots [4,5]. Meiotic DSBs are generated by an endonuclease that comprises SPO11 and TOPOVIBL subunits [2,3,6]. In mice, mutations in *Spo11* result in fewer DSBs during leptotene and zygotene, and defects in the pairing and synapsis of homologous chromosomes [7–9]. The overall amount of SPO11 activity appears to be dynamically controlled at multiple levels during meiotic prophase. At the RNA level, *Spoll* is alternatively spliced into two major isoforms, which appear to have distinct biological functions and whose relative abundance changes as meiotic prophase proceeds [10–12]. There also appears to be regulation of SPO11 activity at the protein level: negative feedback mechanisms acting through the DNA damage-associated protein kinase ATM prevent excessive *Spo11*-dependent DSBs from being generated during meiosis, potentially limiting any genetic instability caused by errors arising during repair of the DSBs and meiotic arrest caused by unrepaired DSBs [13]; whereas positive feedback mechanisms stimulate SPO11 activity in chromosomal regions that remain asynapsed during zygotene, helping to ensure that the homology search is successful and chromosomes synapse fully [14].

Mutations in genes involved in positively regulating SPO11 activity in mammals might be expected to phenocopy *Spo11^-/-^* mutants to some extent in having reduced numbers of DSBs in leptotene, and arrest at pachytene with chromosome asynapsis. One group of genes that is required for chromosome synapsis in mouse spermatocytes, but whose mechanistic role in meiosis is poorly defined, is the germline genome defence genes [15]. These genes are involved in suppressing the activity of retrotransposons in developing germ cells, and mutations in many of them cause defects in progression through the pachytene stage of meiosis [15]. Mutations in one of these germline genome defence genes, *Mael*, which encodes a conserved component of the piRNA pathway, causes de-repression of retrotransposons and a considerable increase in *Spo11*-independent DSBs that are thought to represent DNA damage generated by *de novo* retrotransposition events [16]. In contrast, spermatocytes carrying mutations in the DNA methyltransferase accessory factor *Dnmt3l* also de-repress retrotransposons, but have relatively normal levels of DSBs that are aberrantly distributed across the genome [17–19]. Alternative explanations for the chromosome synapsis defects in genome defence mutants include changes in the meiotic chromatin environment impairing the homology search or synapsis, or physical interactions between the meiotic machinery and either retrotransposon-encoded proteins or genome defence systems affecting meiotic chromosome behaviour [15,20].

In this study we elucidate why loss of the germline genome defence gene *Tex19.1* results in chromosome asynapsis in male meiosis. We show that loss of *Tex19.1* generates a meiotic phenotype distinct from either *Mael^-/-^* or *Dnmt3l^-/-^* mutants. Rather loss of *Tex19.1* phenocopies hypomorphic *Spo11* mutants and impairs DSB formation and the generation of early meiotic recombination foci during the leptotene stage of meiotic prophase. Furthermore, we show that mice carrying mutations in *Ubr2*, which encodes an E3 ubiquitin ligase that physically interacts with TEX19.1, phenocopy the recombination defects seen in leptotene *Tex19.1^-/-^* spermatocytes. These data show that *Tex19.1* and *Ubr2* are required for mice to generate sufficient meiotic DSBs to ensure robust identification and synapsis of homologous chromosomes in meiotic spermatocytes.

## Results

### Chromosome Asynapsis in *Tex19.1^-/-^* Spermatocytes is not Caused by Primary Defects in Synaptonemal Complex Assembly

*Tex19.1* is a DNA methylation-sensitive germline genome defence gene whose expression is primarily restricted to germ cells and pluripotent cells in the embryo [21–24]. We and others have previously reported that *Tex19.1^-/-^* males have defects in spermatogenesis on a mixed genetic background, and that around 50% of pachytene spermatocytes in *Tex19.1^-/-^* testes have asynapsed chromosomes, but the molecular explanation for this defect remains unknown [15,23,25]. Synapsis requires the accurate and timely execution of a number of events in the preceding stages of the first meiotic prophase, including the generation of meiotic DNA double-strand breaks (DSBs) in leptotene, followed by homolog pairing and assembly of the synaptonemal complex (SC) in zygotene [1]. To investigate the molecular basis for the chromosome asynapsis in pachytene *Tex19.1^-/-^* spermatocytes we sought to test whether each of these events occurs normally in the absence of *Tex19.1*.

First, we confirmed that the meiotic chromosome asynapsis phenotype persists in *Tex19.1^-/-^* spermatocytes after backcrossing onto an inbred C57BL/6 genetic background: 65% of *Tex19.1^-/-^* pachytene spermatocytes were asynapsed in this genetic background (Fig 1A), similar to the 50% asynapsis in pachytene previously reported for a mixed genetic background [23,25]. To assess whether the chromosome asynapsis in *Tex19.1^-/-^* spermatocytes represents defects in SC assembly rather than pairing of homologous chromosomes, we scored the configuration of the asynapsed chromosomes in asynapsed pachytene *Tex19.1^-/-^* nuclei. Defects in assembly of the SC transverse filaments results in asynapsed chromosomes that are aligned in their homolog pairs whereas defective recombination or pairing between homologous chromosomes manifests as isolated asynapsed single chromosomes, partial synapsis between non-homologous chromosomes, and incomplete synapsis between homologous chromosomes [7,8,26,27]. Asynapsed chromosomes in *Tex19.1^-/-^* spermatocytes are present in multiple configurations consistent with defects in recombination or homolog pairing, but do not present as asynapsed aligned homolog pairs (Fig 1A, Fig 1B).

**Fig 1.**
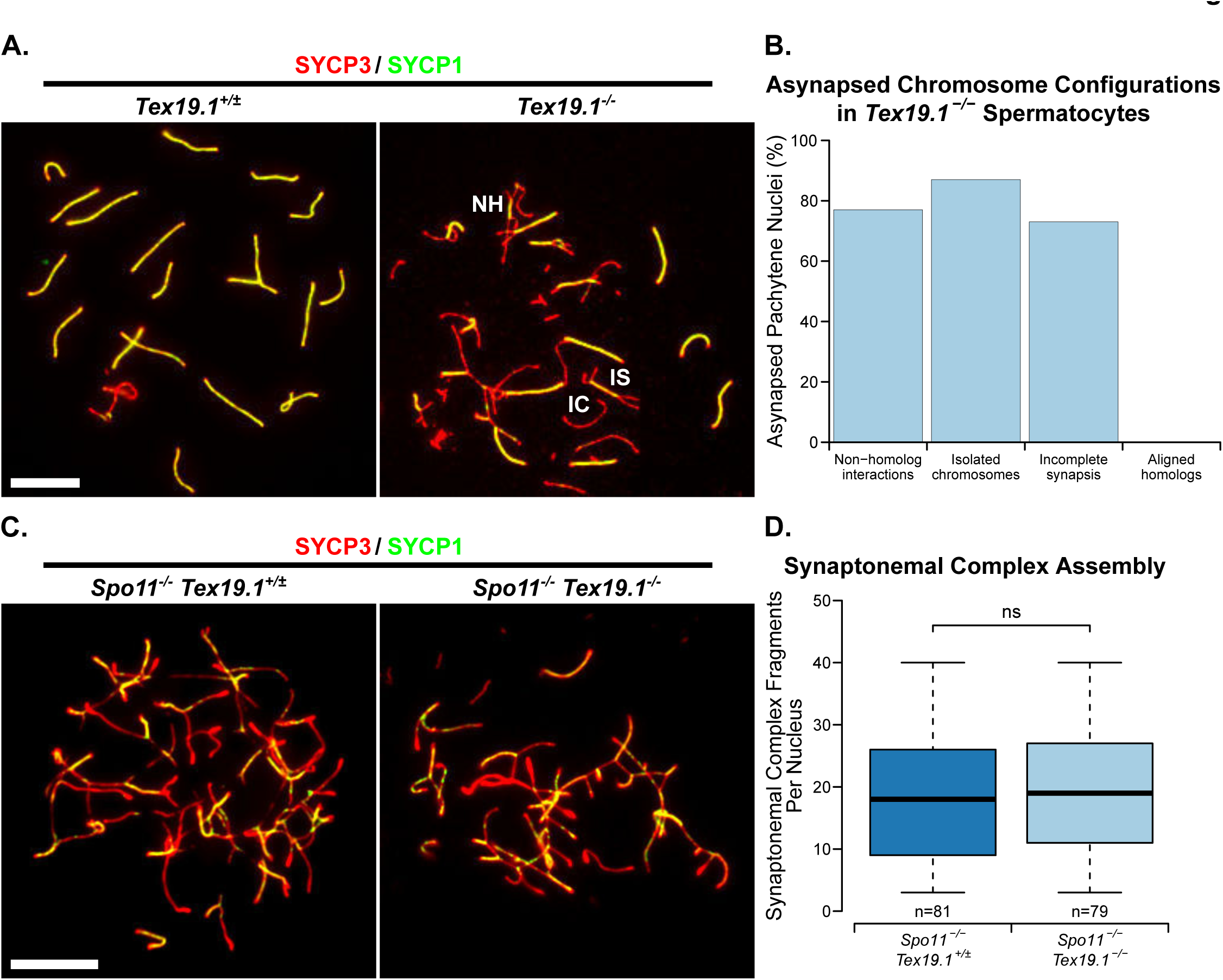
Asynapsis in *Tex19.1^-/-^* Spermatocytes is not Directly Caused by Impaired Synaptonemal Complex Assembly. (A) Immunostaining of chromosome spreads from *Tex19.1*^*+/±*^ and *Tex19.1^-/-^* spermatocytes for synaptonemal complex (SC) components SYCP3 (red) and SYCP1 (green). Asynapsed chromosomes assemble SYCP3 but not SYCP1, and examples involving non-homologous interactions (NH), incomplete synapsis between homologs (IS), and isolated chromosomes (IC) are labelled. Scale bar 10 μm. (B) Percentage of asynapsed pachytene *Tex19.1^-/-^* spermatocytes exhibiting the indicated categories of asynapsed chromosomes (n=60 spreads from 3 mice). Each nucleus is typically represented in more than one category. (C) Immunostaining of chromosome spreads from *Spo11^*-/-*^ Tex19.1^*+/±*^* and *Spo11^-/-^ Tex19.1^-/-^* spermatocytes for the SC components SYCP3 (red) and SYCP1 (green). Fragments of fully assembled SC can be seen where SYCP3 and SYCP1 co-localise. Scale bar 10 μm. (D) Boxplots showing quantification of SC fragments in *Spo11*^*-/-*^*Tex19.1*^*+/±*^and *Spo11^-/-^ Tex19.1^-/-^* spermatocyte nuclei (17.8±1.1 and 19.2±1.1 fragmentsrespectively). n=79, 79 spreads from 3 mice per genotype. ns indicates no significant difference (Mann-Whitney U test).

To confirm that the asynapsis phenotype *Tex19.1^-/-^* spermatocytes does not represent a primary defect in SC assembly, we quantified the effect of *Tex19.1* on the number of SC fragments assembled independently of recombination in a *Spo11^-/-^* genetic background [28]. *Spo11^-/-^ Tex19.1^+/±^* and *Spo11^-/-^ Tex19.1^-/-^* spermatocytes are able to assemble similar amounts of SC in this assay (Fig 1C, Fig 1D), suggesting that loss of *Tex19.1* does not severely impair recombination-independent SC assembly. Taken together, these data suggest that the chromosome asynapsis in *Tex19.1^-/-^* spermatocytes is likely primarily caused by defects in meiotic recombination and/or homolog pairing rather than a direct defect in SC assembly.

### Chromosome Asynapsis in *Tex19.1^-/-^* Spermatocytes is Associated with an Earlier Reduction in the Number of Meiotic Recombination Foci

We next investigated whether loss of *Tex19.1* impaired the generation of recombination intermediates required for homologous chromosome pairing and synapsis. Chromosome spreads were immunostained for the SC components SYCP3 and SYCE2 to identify zygotene nuclei, and for the single-stranded DNA binding proteins RPA, DMC1 and RAD51 to visualise recombination foci associated with the chromosome axes [29]. The number of RPA, DMC1 and RAD51 foci in control *Tex19.1*^*+/±*^ zygotene nuclei (Fig 2) are all within the ranges previously reported for wild-type zygotene spermatocytes (150-250 for RPA foci, 100-250 for DMC1 and RAD51 foci, [29]. However, zygotene *Tex19.1^-/-^* spermatocytes have fewer DMC1 and RAD51 foci than their littermate controls, with DMC1 and RAD51 foci frequency reduced to 87% and 67% of control levels respectively (Fig 2). Interestingly, the number of RPA foci is not statistically different from zygotene control nuclei (Fig 2). The differential behaviour of RPA foci and RAD51 and DMC1 foci in *Tex19.1^-/-^* spermatocytes suggests that the kinetics of recombination foci generation or maturation is perturbed in the absence of *Tex19.1*.

**Fig 2.**
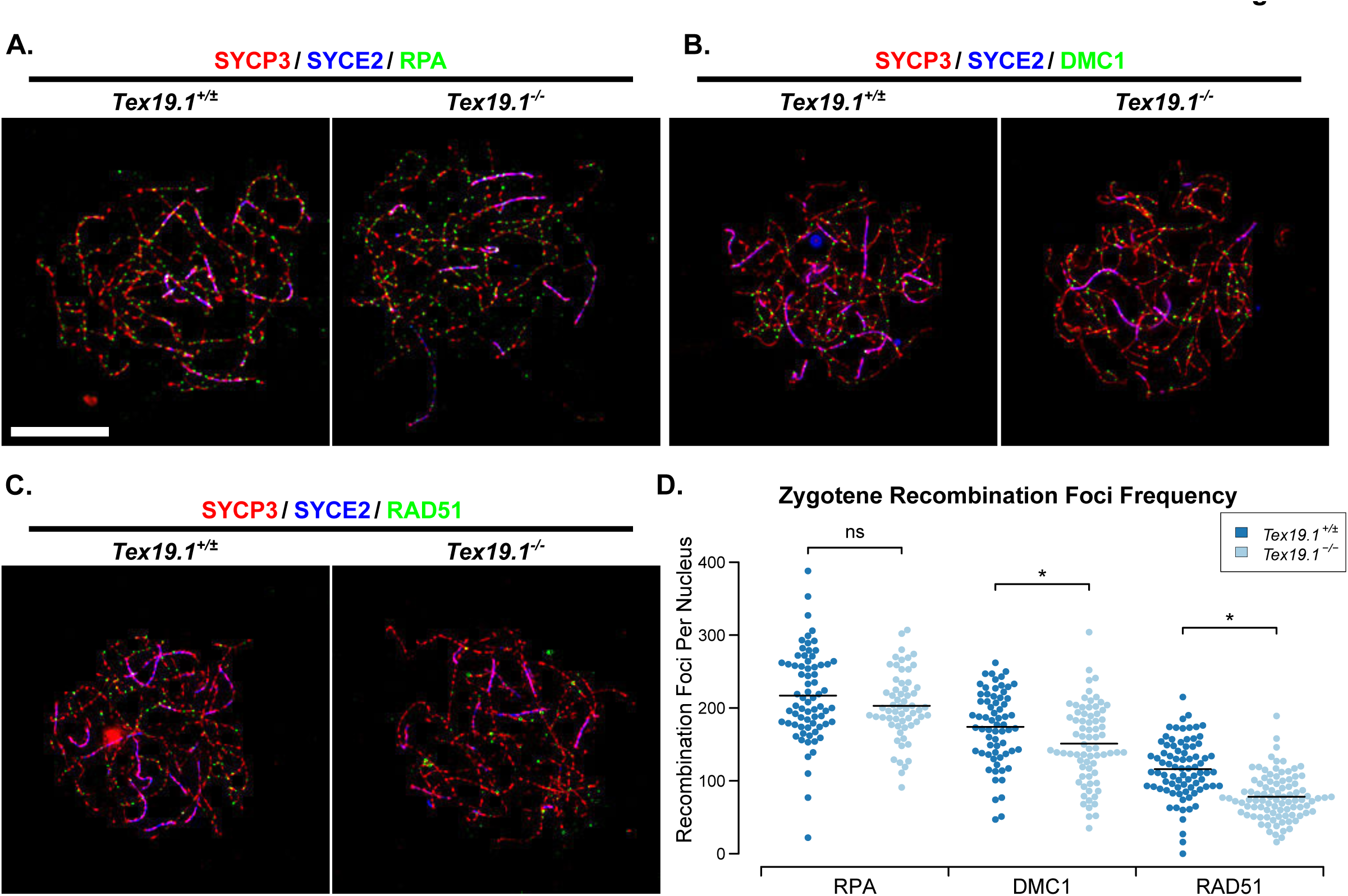
*Tex19.1^-/-^* Spermatocytes Have Reduced Numbers of Recombination Foci During Zygotene. (A-C). Immunostaining of chromosome spreads from *Tex19.1*^*+/±*^ and *Tex19.1^-/-^* spermatocytes for the SC components SYCP3 (red) and SYCE2 (blue) to identify zygotene nuclei and chromosome axes, and RPA (A), DMC1 (B) and RAD51 (C) to mark recombination foci (green). Scale bar 10 μm. (D) Quantification of the number ofRPA, DMC1 and RAD51 recombination foci in zygotene *Tex19.1*^*+/*^and *Tex19.1^-/-^* spermatocytes. n=71, 61, 66, 73, 81, 93 from three mice per genotype. Means are indicated with horizontal bars, * indicates p<0.01, and ns indicates no significant difference (Mann-Whitney U test). Control *Tex19.1*^*+/±*^ zygotene nuclei have 217±7 RPA, 174±6 DMC1, and 116±4 RAD51 foci; *Tex19.1^-/-^* zygotene nuclei have 202±6 RPA, 151±6 DMC1, and 78±3 RAD51 foci.

Meiotic recombination is initiated during leptotene [9] therefore we next investigated whether loss of *Tex19.1* might perturb recombination foci frequency at this earlier stage of meiotic prophase. Counts of RPA, DMC1 and RAD51 foci in leptotene nuclei revealed a severe reduction in the frequency of each of these foci in the absence of *Tex19.1* (Fig 3). The numbers of RPA foci, DMC1 foci and RAD51 foci in leptotene *Tex19.1^-/-^* spermatocytes were reduced to 63%, 30%, and 60% of those present in control spermatocytes (Fig 3). Thus, *Tex19.1* is required to generate the correct number of meiotic recombination foci in leptotene spermatocytes, a defect that precedes the chromosome asynapsis at pachytene.

**Fig 3.**
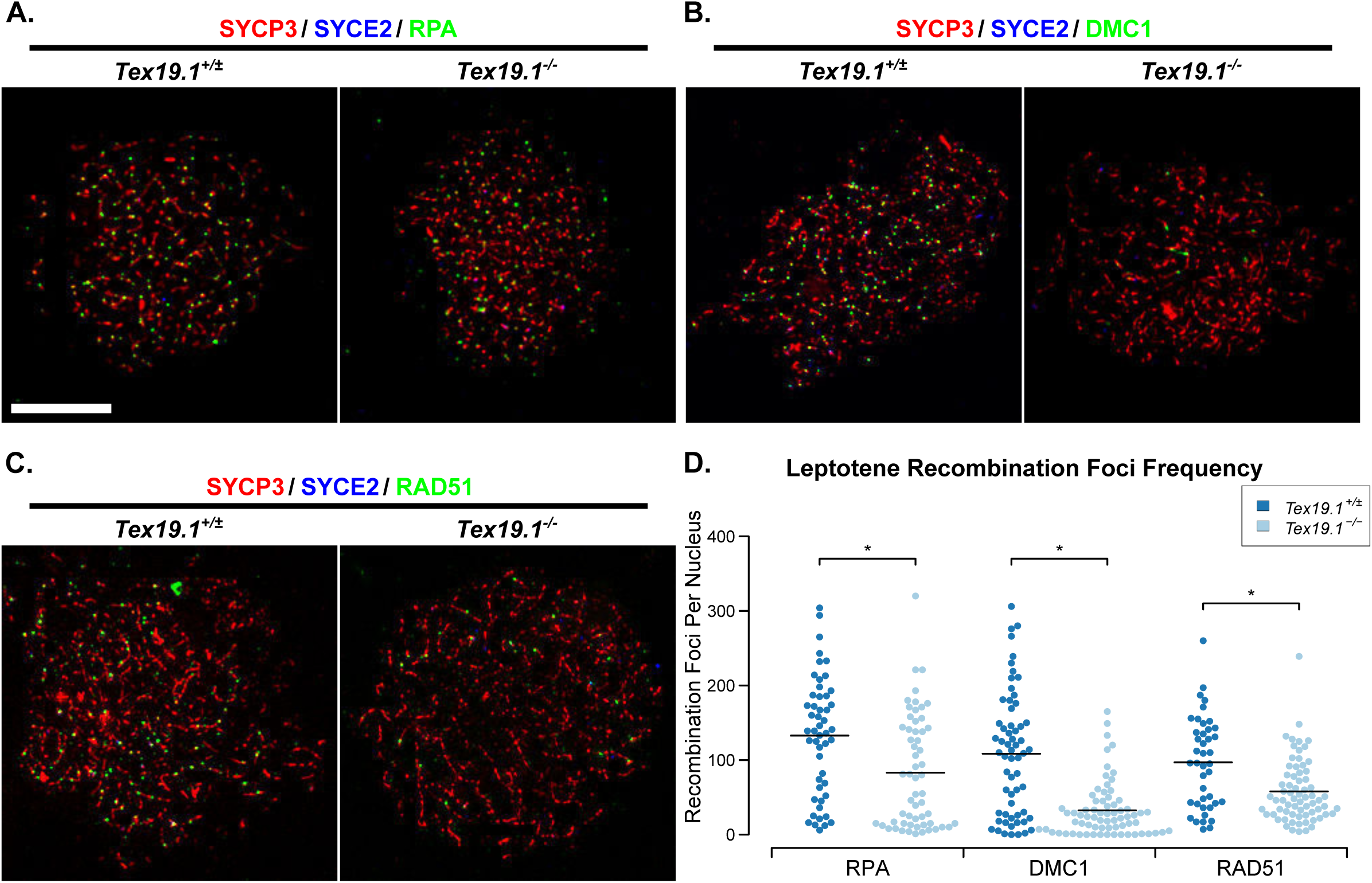
*Tex19.1^-/-^* Spermatocytes Have Reduced Numbers of Recombination Foci During Leptotene. (A-C) Immunostaining of chromosome spreads from *Tex19.1*^*+/±*^ and *Tex19.1^-/-^* spermatocytes for the SC components SYCP3 (red) and SYCE2 (blue) to identify leptotene nuclei and fragments of chromosome axes, and RPA (A), DMC1 (B), and RAD51 (C) to mark recombination foci (green). Scale bar 10 μm. (D) Quantification of the number of RPA, DMC1 and RAD51-positive recombination foci in leptotene *Tex19.1*^*+/±*^ and *Tex19.1^-/-^* spermatocytes. n=50, 58, 61, 71, 42, 69 from three mice per genotype. Means are indicated with horizontal bars, and * indicates p<0.01 (Mann-Whitney U test). Control *Tex19.1*^*+/±*^ leptotene nuclei have 133±11 RPA, 108±11 DMC1, and 97±9 RAD51 foci; *Tex19.1^-/-^* leptotene nuclei have 83±10 RPA, 32±4 DMC1, and 58±5 RAD51 foci.

The reduced numbers of recombination foci in *Tex19.1^-/-^* spermatocytes could potentially decrease the efficiency of the DSB-dependent homology search and contribute to chromosome asynapsis in this mutant. Analysis of *Spo11* hypomorphs suggests that reduced numbers of meiotic DSBs impairs the initiation of synapsis and manifests as reduced numbers of synaptonemal complex fragments during late leptotene/early zygotene stages. We therefore analysed the extent of synapsis in zygotene *Tex19.1^-/-^* nuclei to assess whether the initiation of synapsis might similarly be impaired in these mutants. Chromosome spreads were immunostained with axial and central element synaptonemal complex markers and the percentage synapsis assessed in each zygotene nucleus (Supporting Fig S1). In the absence of *Tex19.1*, most zygotene nuclei contained very low amounts of synapsis (<10%), whereas the majority of control zygotene nuclei contained intermediate levels of synapsis (10-70%, Supporting Fig S1). The extent of synapsis in *Tex19.1^-/-^* nuclei is more consistent with these mutants exhibiting a widespread block or delay in the initiation of synapsis throughout the nucleus, rather than defects in synapsis of specific chromosomes or progression of synapsis along the chromosome axes once it has initiated. Thus, as described for *Spo11* hypomorphs [14], the reduced numbers of DSBs generated in *Tex19.1^-/-^* sperrmatocytes during leptotene could potentially cause defects in homologous chromosome synapsis during zygotene resulting in asynapsis persisting in pachytene.

### *Tex19.1^-/-^* Spermatocytes Have Defects in the Generation of *Spo11*-Dependent Meiotic DSBs

The reduced number of RPA, DMC1 and RAD51 foci in leptotene *Tex19.1^-/-^* spermatocytes might reflect an earlier defect in the formation of DSBs or their subsequent resection to form single-stranded DNA ends. Phosphorylation of the histone variant H2AX to generate γH2AX occurs in response to *Spo11*-dependent DSB formation [9], but is not impaired in spermatocytes defective in the DNA polymerase β-dependent steps that process those DSBs [30]. We therefore tested whether loss of *Tex19.1* affects γH2AX abundance in leptotene spermatocytes. In both control and *Tex19.1^-/-^* leptotene nuclei, γH2AX is present as a diffuse cloud of staining over regions of the nucleus (Fig 4A). Interestingly, quantification of the γH2AX signal showed that the amount of γH2AX in leptotene *Tex19.1^-/-^* nuclei was around half that in *Tex19.1*^*+/±*^ controls (Fig 4B). Taken together, the reduced numbers of recombination foci and the reduced intensity of γH2AX immunostaining in *Tex19.1^-/-^* spermatocytes likely reflects an impaired ability to generate *Spo11*-dependent DSBs.

**Fig 4.**
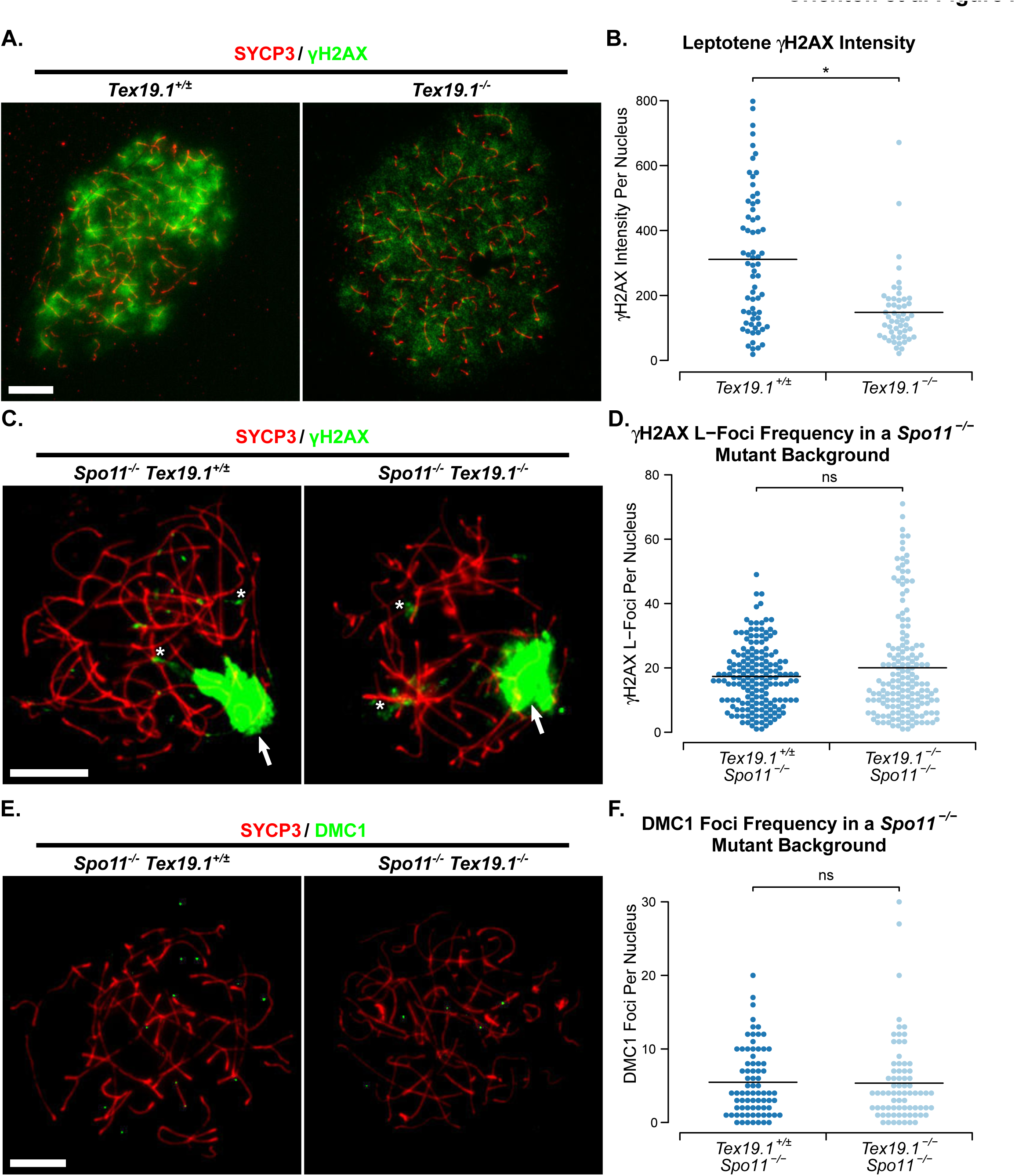
Loss of *Tex19.1* Impairs the Generation of SPO11-Dependent DSBs in Spermatocytes. (A) Immunostaining of chromosome spreads from *Tex19.1*^*+/±*^ and *Tex19.1^-/-^* spermatocytes for the SC component SYCP3 (red) to identify leptotene nuclei and fragments of chromosome axes, and γH2AX (green) as a marker for DSBs. Scale bar 10 μm. (B) Quantification of γH2AX immunostaining intensity (arbitrary units) in leptotene *Tex19.1*^*+/±*^ and *Tex19.1^-/-^* spermatocytes (311±15 and 148±26 units respectively). n=66, 52 from three mice per genotype. Means are indicated with horizontal bars, and * indicates p<0.01 (Mann-Whitney U test). (C, E). Immunostaining of chromosome spreads from *Spo11^-/-^Tex19.1^+/±^* and *Spo11^-/-^ Tex19.1^-/-^* spermatocytes for the SC component SYCP3 (red) to identify chromosome axes, and γH2AX (C, green) or DMC1 (E, green) as markers for DSBs and recombination foci respectively. Arrows in C label the pseudo sex body, asterisks label example axis-associated L-foci. Scale bars 10 μm. D, F. Quantification of γH2AX L-foci (D) and DMC1-positive recombination foci (F) in *Spo11^-/-^Tex19.1^+/±^* and *Spo11^-/-^ Tex19.1^-/-^* spermatocytes. n=169, 162 for D, and 76, 74 for F. Both analyses were performed on spreads from three mice per genotype. Means are indicated with horizontal bars, and ns indicates no significant difference (Mann-Whitney U test). *Spo11*^*-/-*^*Tex19.1*^*+/±*^ and*Spo11^-/-^ Tex19.1^-/-^* spermatocytes have 17.3±0.8 and 20.0±1.3 γH2AX L-foci and 5.5±0.5 and 5.4±0.7DMC1 foci respectively.

The bulk of the γH2AX generated in spermatocytes reflects the generation of *Spo11*-dependent meiotic DSBs, however small amounts of γH2AX are generated independently of *Spo11* in these cells [9,31,32]. The extent of the decrease in γH2AX abundance in *Tex19.1^-/-^* spermatocytes is arguably more consistent with defects in the generation of *Spo11*-dependent DSBs, but it is possible that loss of *Tex19.1* also affects *Spo11*-independent γH2AX generated during leptotene. To test directly whether loss of *Tex19.1* affects *Spo11*-independent γH2AX we quantified γH2AX and DMC1 foci in *Spo11^-/-^ Tex19.1^-/-^* double mutant spermatocytes. The relatively low levels of γH2AX present in *Spo11^-/-^* spermatocytes typically manifests as a large cloud of pseudo sex body staining with smaller additional flares of chromosome axis-associated L-foci [9,31,32]. *Spo11^-/-^ Tex19.1^-/-^* spermatocytes displayed similar γH2AX staining patterns and similar numbers of γH2AX L-foci as *Spo11^-/-^Tex19.1^+/±^* controls (Fig 4C, Fig 4D). In addition, although loss of *Tex19.1* impairs DMC1 foci frequency in a wild-type *Spo11* background (Fig 2, Fig 3), loss of *Tex19.1* has no detectable effect on DMC1 foci frequency in a *Spo11*^*-/-*^ mutant background (Fig 4E, Fig 4F). Thus, loss of *Tex19.1* appears to perturb the generation of *Spo11*-dependent, but not *Spo11*-independent, DSBs in spermatocytes, and in this respect the *Tex19.1^-/-^* phenotype bears some resemblance to hypomorphic *Spo11* mutants [14,33]

SPO11 is locally regulated in the nucleus, and feedback controls are thought to allow SPO11 to continue to generate DSBs on asynapsed regions of the chromosomes in late zygotene [14]. *Spo11* hypomorphs are still able to act on asynapsed chromatin to elevate DSB frequency in these regions of asynapsis [14]. To assess whether asynapsed chromatin is similarly able to accumulate high levels of DSBs in *Tex19.1^-/-^* mutants, we counted the number of recombination foci associated with the sex chromosomes, which remain largely asynapsed during pachytene. In the absence of *Tex19.1*, sex chomosomes were still able to accumulate similar numbers of DSBs during pachytene as in control pachytene nuclei (Supporting Fig S1). Thus, like *Spo11* hypomorphs, loss of *Tex19.1* does not prevent the accumulation of DSBs on asynapsed chromatin.

### Loss of *Tex19.1* Does Not Impair MEI4 Localisation or H3K4me3 Deposition at Recombination Hotspots

We next investigated whether the impaired generation of *Spo11*-dependent leptotene DSBs in *Tex19.1^-/-^* spermatocytes might reflect defects upstream of *Spo11* in meiotic recombination. The requirements upstream of *Spo11* for meiotic DSB formation are relatively poorly understood in mammals, however *Spo11* does depend on the recruitment of the conserved axis-associated protein MEI4 to the chromosomal axes in leptotene [34]. We therefore quantified MEI4 foci in leptotene *Tex19.1^-/-^* nuclei to test whether this early requirement for meiotic DSB generation is perturbed by loss of *Tex19.1*. Control leptotene *Tex19.1*^*+/±*^ spermatocytes possess an average of 218 axis-associated MEI4 foci (Fig 5A, Fig 5B), similar but slightly lower than the average 309 foci per leptotene nucleus reported previously [34]. Leptotene *Tex19.1^-/-^* nuclei possess similar numbers of MEI4 foci as leptotene *Tex19.1*^*+/±*^ controls (Fig 5A, Fig 5B). Thus, the reduced meiotic DSB frequency seen in *Tex19.1^-/-^* spermatocytes appears to be a consequence of defects acting downstream or independently of MEI4 localisation to chromosome axes.

**Fig 5.**
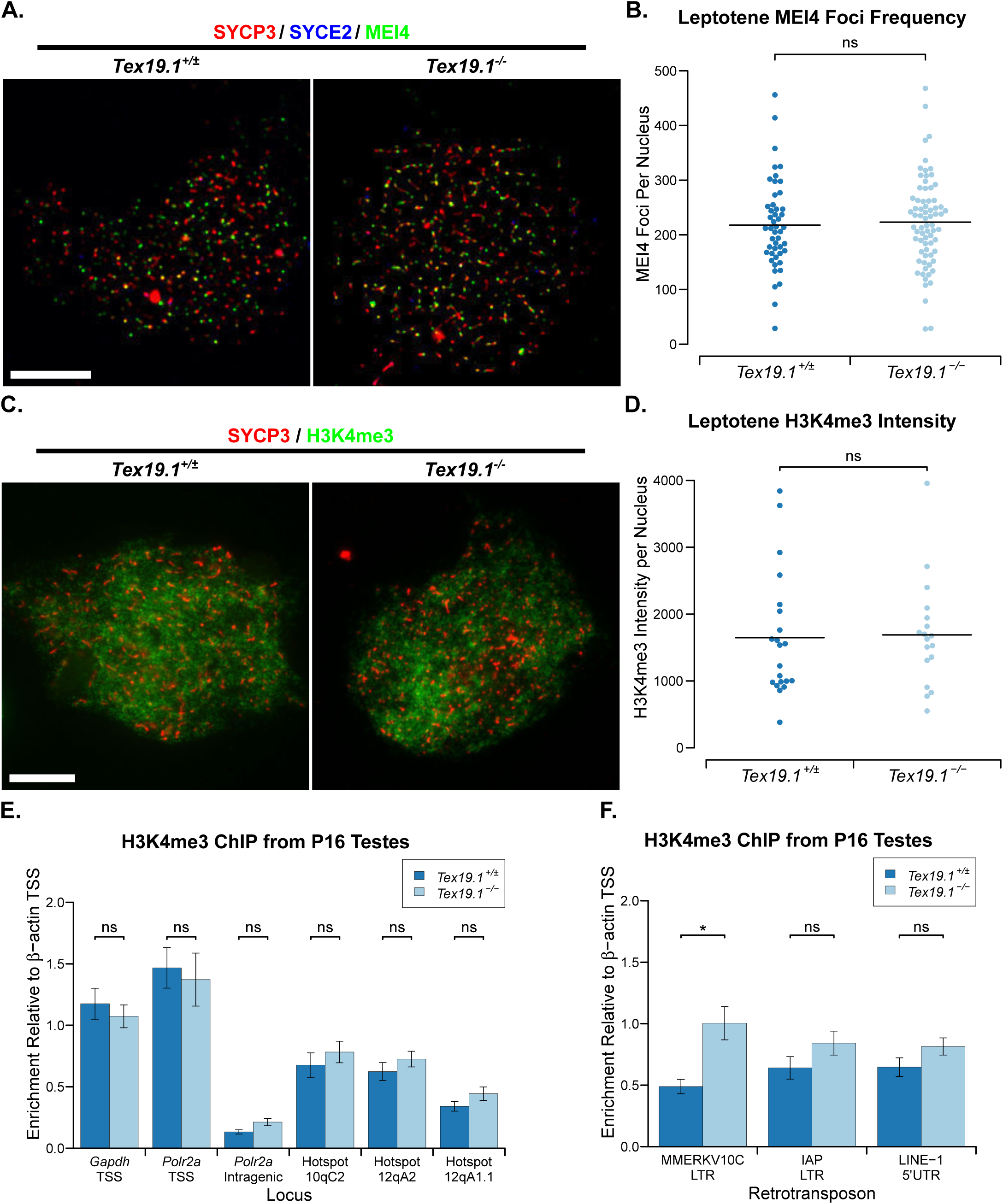
*Tex19.1^-/-^* Spermatocytes Have No Overt Defects in MEI4 Localisation or in H3K4me3 Accumulation at Recombination Hotspots. (A) Immunostaining of spermatocyte chromosome spreads for the SC components SYCP3 (red) to identify leptotene nuclei and fragments of chromosome axes, and MEI4 (green). Scale bar 10 μm. Quantification of MEI4 foci in leptotene spermatocytes (218±12 for *Tex19.1*^*+/±*^, 223±10 for *Tex19.1^-/-^*, n=48, 72 from three mice per genotype). Means are indicated with horizontal bars, ns indicates no significant difference (Mann-Whitney U test). (C) Immunostaining of spermatocyte chromosome spreads for the SC components SYCP3 (red) and SYCE2 (blue) to identify leptotene nuclei, and H3K4me3 (green). Scale bar 10 μm. (D) Quantification of anti-H3K4me3 staining intensity (1648±202 and 1689±187 arbitrary units respectively, n=21, 18 from three mice per genotype). Means are indicated with horizontal bars, ns indicates no significant difference (Mann-Whitney U test). (E, F) H3K4me3 chromatin immunoprecipitation (ChIP) from P16 *Tex19.1*^*+/±*^ and *Tex19.1^-/-^* testes. qPCR for recombination hotspots (E) and retrotransposon sequences (F) was performed on H3K4me3 ChIP and abundance measured relative to input chromatin, then normalised to enrichment for the β-actin (*Actb*) transcriptional start site (TSS). Mean normalised enrichment ± standard error from three animals of each genotype is shown. *Polr2a* and *Gapdh* TSSswere used as positive controls, and an intragenic region of *Polr2a* as a negative control. ns indicates no significant difference, * indicates p<0.05 (Student’s t-test).

*Spo11* function is also influenced by the activity of the histone methyltransferase PRDM9, which targets SPO11 to recombination hotspots [2,4,5]. Mutations in *Prdm9* result in reduced anti-H3K4me3 immunostaining in P14 spermatocytes, a failure to enrich H3K4me3 at *Prdm9*-dependent recombination hotspots, a reduction in recombination foci during early prophase, and meiotic chromosome asynapsis [5,35,36]. We therefore tested whether loss of *Tex19.1* might impair *Prdm9* function by assessing anti-H3K4me3 immunostaining intensity in leptotene nuclei. However, we could not detect a difference in the amount of anti-H3K4me3 immunostaining between *Tex19.1*^*+/±*^and *Tex19.1^-/-^* leptotene nuclei (Fig 5C, Fig 5D). To test whether the distribution of H3K4me3 rather than its total abundance might be altered in the absence of *Tex19.1* we performed H3K4me3 chromatin immunoprecipitation (ChIP) on P16 testes. H3K4me3 is enriched at transcriptional start sites (TSSs) of active genes in addition to meiotic recombination hotspots [37], and as expected both *Tex19.1*^*+/±*^ and *Tex19.1^-/-^* testes show enrichment of H3K4me3 at *Gapdh* and *Polr2a* active TSSs, but not at a *Polr2a* intragenic region (Fig 5E). However, loss of *Tex19.1* does not perturb the accumulation of H3K4me3 at *Prdm9*-dependent recombination hotspots (Fig 5E). Thus, the defects in *Spo11*-dependent DSB formation seen in *Tex19.1^-/-^* spermatocytes does not appear to be a downstream consequence of impaired *Prdm9* activity.

*Tex19.1* plays a role in repressing retrotransposons in testes and placenta [23,38,39], and *Tex19.1^-/-^* testes have increased abundance of *MMERVK10C* retrotransposon RNA, but not RNAs encoding *IAP* or *LINE-1* retrotransposons [23,38]. To test if the increase in *MMERVK10C* RNA is a consequence of transcriptional de-repression we also analysed retrotransposon sequences in the P16 testis H3K4me3 ChIP. Interestingly, the LTR driving *MMERVK10C* expression, but not *IAP* LTRs or *LINE-1* 5' UTR sequences are enriched in anti-H3K4me3 ChIP from *Tex19.1^-/-^* testes relative to *Tex19.1*^*+/±*^ controls (Fig 5F). Thus the increase in *MMERVK10C* retrotransposon RNA abundance previously reported in *Tex19.1^-/-^* testes [23,38] reflects, at least in part, transcriptional de-repression of this element. However, the 2-fold increase in H3K4me3 abundance at *MMERVK10C* LTR sequences does not detectably interfere or compete with enrichment of H3K4me3 at *Prdm9*-dependent recombination hotspots.

### The *Tex19.1^-/-^* Meiotic Recombination Defect is Phenocopied by Mutations in *Ubr2*

*Tex19.1* encodes a protein of unknown biochemical function that physically interacts with the E3 ubiquitin ligase UBR2 [25]. TEX19.1 protein is undetectable in *Ubr2*^*-/-*^ testes, suggesting that much of the TEX19.1 protein in the testis requires UBR2 for its stability [25]. *Ubr2* is implicated in the ubiquitylation and degradation of N-end rule substrates and previous reports suggest that loss of *Ubr2* causes variable defects in spermatogenesis possibly depending on the strain background [40]. Some *Ubr2*^*-/-*^ spermatocytes are reported to progress into pachytene and arrest due to defects in the accumulation of ubiquitylated histone H2A at the sex body and meiotic sex chromatin inactivation during pachytene [41,42]. *Ubr2*^*-/-*^ spermatocytes are also reported to arrest and apoptose in prophase I due to defects in the repair of DSBs, homologous chromosome pairing, and synaptonemal complex formation [40,42]. Given the lack of detectable TEX19.1 protein in *Ubr2*^*-/-*^ testes, we tested whether the reported defects in homologous chromosome pairing and synaptonemal complex formation in *Ubr2*^*-/-*^ spermatocytes [40] might reflect earlier defects in the initiation of meiotic recombination similar to *Tex19.1^-/-^* spermatocytes. We generated *Ubr2*^*-/-*^ mice carrying a premature stop codon in the N-terminal region of UBR2 within the UBR domain that binds N-end rule substrates. The *Ubr2*^*-/-*^ mice analysed here have no detectable UBR2 protein in their testes (Supporting Fig S2A), a 68% reduction in testis weight (Supporting Fig S2B), and no detectable sperm in their epididymis (Supporting Fig S2C), consistent with *Ubr2*^*-/-*^ spermatogenesis defects reported previously [40]. The seminiferous tubules in *Ubr2*^*-/-*^ mice contain reduced numbers of post-meiotic round and elongated spermatids, accumulations of pyknotic and zygotene-like nuclei consistent with meiotic defects (Supporting Fig S2D) as reported previously [40,42]. Similar to *Tex19.1^-/-^* testes [23], some round and elongated post-meiotic spermatids are detectable in *Ubr2*^*-/-*^testes suggesting that any meiotic defects present do not completely block spermatogenesis. Furthermore, loss of *Ubr2* phenocopies the specific retrotransposon derepression seen in *Tex19.1^-/-^* testes [23]: *MMERVK10C*, but not *LINE-1* or *IAP*, retrotransposon RNAs are derepressed in *Ubr2*^*-/-*^spermatocytes (Fig 6A, Fig 6B).

**Fig 6.**
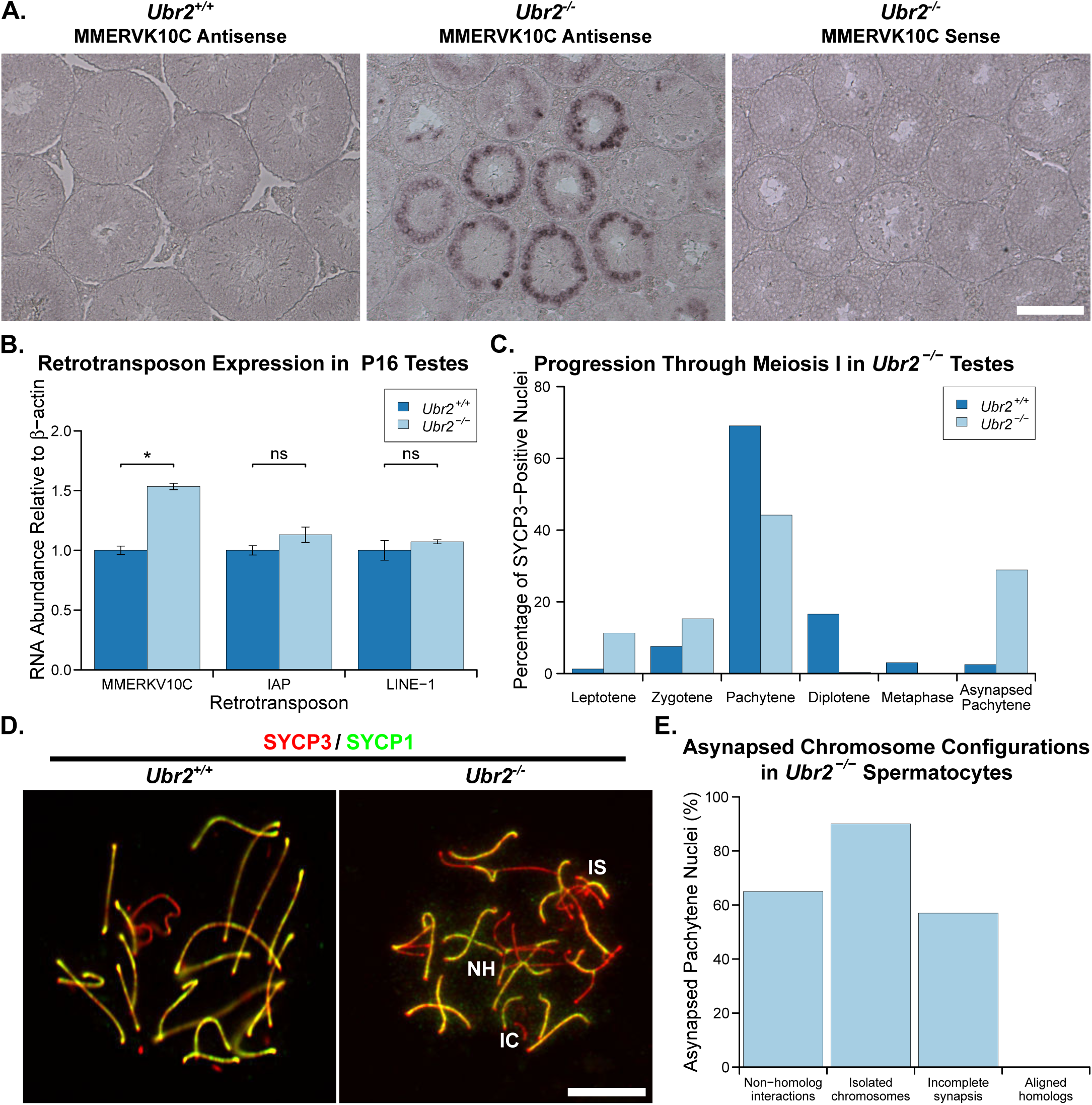
*Ubr2*^*-/-*^Spermatocytes Phenocopy the Retrotransposon Derepression and Asynapsis Phenotypes Present in *Tex19.1^-/-^* Mutants. (A) In situ hybridisation for *MMERVK10C* retrotransposon RNA in *Ubr2*^*-/-*^ testes. Specific signal (dark purple precipitate) is present in *Ubr2*^*-/-*^ spermatocytes hybridised with an antisense *MMERVK10C* probe. Scale bar 100 μm. (B) qRT-PCR for *MMERVK10C*, *IAP* and *LINE-1* retrotransposons in P16 *Ubr2*^*-/-*^ testes. Mean abundance of retrotransposon RNAs relative to β-actin is shown for two *Ubr2*^*+/+*^ and three *Ubr2*^*-/-*^ animals. ns indicates no significant difference, * indicates p<0.05 (Student’s t-test). (C) Chromosome spreads from testes from three *Ubr2*^*+/+*^ and three *Ubr2*^*-/-*^ animals were immunostained with antibodies to SYCP3 and SYCP1, and SYCP3-positive nuclei scored for meiotic substage (n=398, 595). *Ubr2*^*-/-*^ spreads contained a significant proportion of aberrant pachyene nuclei containing asynapsed chromosomes, but few diplotene or metaphase I nuclei compared to *Ubr2*^*+/+*^ controls. (D) Immunostaining of chromosome spreads from *Ubr2*^*+/+*^and *Ubr2*^*-/-*^spermatocytes for SYCP3 (red) and SYCP1 (green). Asynapsed chromosomes assemble SYCP3 but not SYCP1, and examples involving non-homologous interactions (NH), incomplete synapsis between homologs (IS), and isolated chromosomes (IC) are labelled. Scale bar 10 μm. (E) Percentage of asynapsed pachytene *Ubr2*^*-/-*^ spermatocytes exhibiting the indicated categories of asynapsed chromosomes (n=101 from a total of three mice). Each nucleus is typically represented in more than one category.

We tested whether the meiotic defects in *Ubr2*^*-/-*^ spermatocytes might resemble the asynapsis seen in *Tex19.1^-/-^* spermatocytes (Fig 1A, 1B). Chromosome spreads from *Ubr2*^*-/-*^ testes confirm that this *Ubr2* mutant allele causes defects in progression through meiotic prophase, and very few spermatocytes progress through pachytene into diplotene (Fig 6C). Furthermore, around 40% of pachytene *Ubr2*^*-/-*^ spermatocytes had at least one asynapsed autosome pair when staging SYCP3-positive nuclei for meiotic progression under low magnification (Fig 6C, Fig 6D). At higher magnification around 65% *Ubr2*^*-/-*^ pachytene nuclei have some autosomal asynapsis. Like in *Tex19.1^-/-^* spermatocytes (Fig 1A, Fig 1B), these asynapsed chromosomes are present in multiple configurations consistent with defects in recombination or homolog pairing (Fig 6E). Similar to *Tex19.1^-/-^* spermatocytes, the asynapsis in *Ubr2*^*-/-*^ spermatocytes is also associated with earlier defects in the generation of meiotic DSBs. γH2AX abundance and DMC1 foci frequency are reduced to around 50% and 52% respectively of those seen during leptotene in *Ubr2*^*-/-*^ mutants (Fig 7), which contrasts with a previous report that γH2AX staining, and RAD51 and RPA foci frequency are unaffected in leptotene *Ubr2*^*-/-*^ spermatocytes [42]. Consistent with the decrease in the generation of meiotic DSBs during leptotene reported here, DMC1 foci frequency remains around 66% of that seen in control spermatocytes during zygotene (Fig 7). Despite the qualitative similarity between the defects in meiotic DSB formation in *Ubr2*^*-/-*^ and *Tex19.1^-/-^* spermatocytes, the extent of the recombination defect in *Ubr2*^*-/-*^ spermatocytes appears to be slightly less than that in *Tex19.1^-/-^* spermatocytes at leptotene. However, both these mutants have similar reductions in DMC1 foci counts in zygotene. We therefore tested whether the reduced meiotic DSB formation in *Ubr2*^*-/-*^spermatocytes would be sufficient to delay or impair the initiation of chromosome synapsis as described for *Tex19.1^-/-^* spermatocytes (Supporting Fig S1) and for *Spo11* hypomorphs [14]. Measurement of the extent of chromosome synapsis in zygotene *Ubr2*^*-/-*^ spermatocytes suggests that, like in *Tex19.1^-/-^* mutants and *Spo11* hypomorphs, synapsis is delayed in the absence of *Ubr2*. (Supporting Fig S2D, Supporting Fig S2E). These data suggest that the defect in progression to pachytene previously reported in *Ubr2*^*-/-*^ mutants [40] may reflect loss of TEX19.1 protein and earlier defects in the initiation of recombination in these mutants. Furthermore, these data show that *Ubr2* and *Tex19.1* are both required to generate sufficient meiotic DSBs to drive robust homologous chromosome synapsis in mouse spermatocytes.

**Fig 7.**
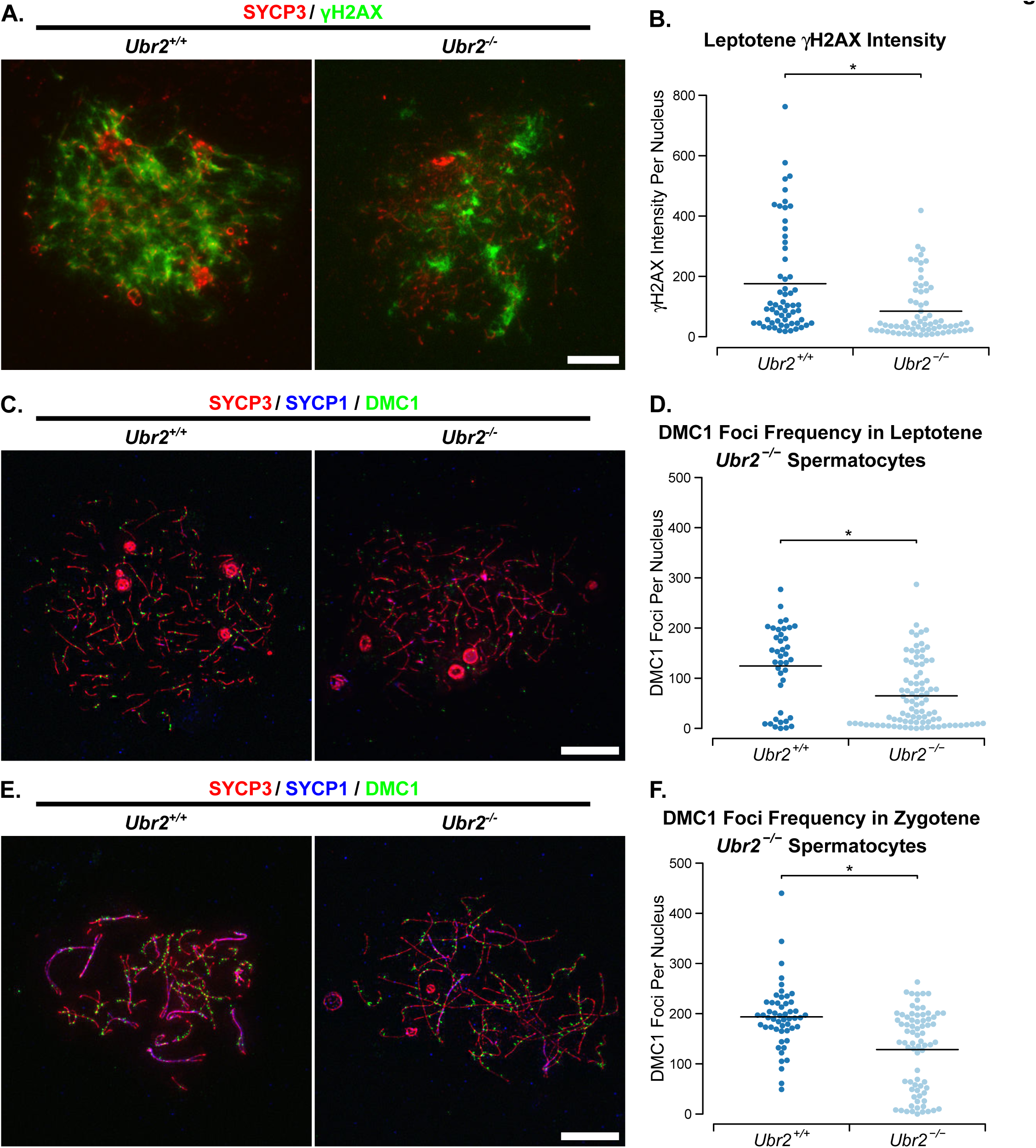
*Ubr2*^*-/-*^Spermatocytes Have Defects in the Initiation of Meiotic Recombination. (A) Immunostaining of chromosome spreads from *Ubr2*^*+/+*^ and *Ubr2*^*-/-*^ spermatocytes for the SC component SYCP3 (red) to identify leptotene nuclei and fragments of chromosome axes, and γH2AX (green) as a marker for DSBs. Scale bar 10 μm. (B) Quantification of γH2AX immunostaining intensity (arbitrary units) in leptotene *Ubr2*^*+/+*^ and *Ubr2*^*-/-*^ spermatocytes (176±23 and 85±11 units respectively). n=60, 68 from three mice per genotype. Means are indicated with horizontal bars, and * indicates p<0.01 (Mann-Whitney U test). (C, E) Immunostaining of chromosome spreads from *Ubr2*^*+/+*^ and *Ubr2*^*-/-*^ spermatocytes for the SC components SYCP3 (red) and SYCP1 (blue) to identify leptotene nuclei (C), zygotene nuclei (E), and fragments of chromosome axes; and DMC1 (green) to mark recombination foci. Scale bar 10 μm. (D, F) Quantification of the number of DMC1-positive recombination foci in spermatocytes from two *Ubr2*^*+/+*^and three *Ubr2*^*-/-*^spermatocytes during leptotene (D) and zygotene (F). Means are indicated with horizontal bars, and * indicates p<0.05 (Mann-Whitney U test). Control *Ubr2*^*+/+*^ nuclei have 124±13 DMC1 foci in leptotene (n=40) and 194±9 in zygotene (n=52); *Ubr2*^*-/-*^ nuclei have 65±7 DMC1 foci in leptotene (n=89) and 128±9 in zygotene (n=75).

## Discussion

### DSB Frequency and Chromosome Synapsis in Meiosis

This study aimed to elucidate the mechanistic basis of the chromosome synapsis defect in male mice carrying mutations in the germline genome defence gene *Tex19.1* [23]. We have shown that the pachytene chromosome asynapsis in these mice, and in mice carrying mutations in the TEX19.1-interacting protein UBR2, is likely a downstream consequence of reduced numbers of meiotic DSBs earlier in meiotic prophase. Wild-type mice generate around 10-fold more meiotic DSBs than there are chiasmata, and the large numbers of DSBs generated in leptotene and zygotene appear to be important to drive pairing and synapsis of homologous chromosomes [2,7,8]. Allelic series of *Spo11* activity suggest that reducing the number of meiotic DSBs to around 50% of normal levels is sufficient to cause chromosome asynapsis [14,33]. The reduction in meiotic DSBs seen in leptotene *Tex19.1^-/-^* and leptotene *Ubr2*^*-/-*^ spermatocytes, is similar to this threshold and could be sufficient to account for the chromosome asynapsis seen in these mutants. Notably, *Tex19.1^-/-^* spermatocytes and *Ubr2*^*-/-*^ spermatocytes do not exhibit a severe asynapsis phenotype, indeed only a proportion of pachytene *Tex19.1^-/-^* or *Ubr2*^*-/-*^ spermatocytes have asynapsed chromosomes, and there is some progression to post-meiotic spermatid stages in both these mutants. Thus the ~50% reduction in leptotene DSB frequency caused by loss of *Tex19.1* or *Ubr2* could be sufficient to cause the level of asynapsis present in these spermatocytes.

Interestingly, the level of meiotic DSBs in *Tex19.1^-/-^* spermatocytes during zygotene is closer to wild-type levels than that seen during leptotene, suggesting additional DSBs are generated during zygotene that allow the *Tex19.1^-/-^* spermatocytes to catch up with wild-type cells. It is possible that DSB generation is delayed in *Tex19.1^-/-^* spermatocytes, or alternatively this compensation of the *Tex19.1* DSB deficiency during zygotene reflect control mechanisms that regulate DSB frequency in meiotic cells. An overall delay in germ cell development is probably not causing a delay in DSB formation relative to axial element assembly as previous analysis of gene expression profiles in P16 *Tex19.1^-/-^* testes does not exhibit enrichment of genes expressed in more immature germ cells such as spermatogonia or leptotene spermatocytes [23,38]. In hypomorphic *Spo11* mice, additional compensatory DSBs are generated on asynapsed regions of the chromosomes, potentially stimulating homology search and synapsis in these regions [14]. However, although this feedback mechanism might be rescuing asynapsis to some degree, the additional DSBs that are generated during zygotene are not sufficient to allow the majority of *Tex19.1^-/-^* spermatocytes to complete synapsis.

### Meiotic Defects in Genome Defence Mutants

*Tex19.1* is one of a group of germline genome defence genes which cause retrotransposon de-repression and defects in meiotic chromosome synapsis [15]. Although a common mechanism could link de-repression of retrotransposons and chromosome asynapsis in these mutants, mutations in different germline genome defence genes seem to have distinct effects on DNA damage and DSBs during meiosis. Mutations in *Mael* cause a striking increase in *Spo11*-independent DNA damage in meiotic spermatocytes which is thought to represent DSBs generated during retrotransposition [16], but the absence of any detectable increase in *Spo11*-independent DSBs in *Spo11*^*-/-*^*Tex19.1^-/-^* spermatocytes reported here contrasts markedly with the phenotype of *Spo11*^*-/-*^*Mael*^*-/-*^spermatocytes. Moreover, zygotene DSB frequency is reduced in *Tex19.1^-/-^* spermatocytes, but are not perturbed by mutations in *Dnmt3l* [18,19]. Thus, there are phenotypic differences between the meiotic defects in different germline genome defence mutants, and distinct mechanisms may be causing asynapsis in each of these mutants.

The spectrum of retrotransposons de-repressed in *Tex19.1^-/-^* spermatocytes differs from those de-repressed in *Mael*^*-/-*^ testes and *Dnmt3l*^*-/-*^ testes [16,17,23,38]. It is therefore possible that some of the differences between the meiotic phenotypes of these mutants reflects differences in the type of retrotransposon de-repressed or the mechanism of de-repression. Data from *Dnmt3l*^*-/-*^ mice suggests that transcriptional activation of *LINE-1* retrotransposons alters the distribution of meiotic DSBs and induces recombination at *LINE-1* elements leading to interactions between non-homologous chromosomes [19]. It is not clear if transcriptional activation of *MMERVK10C* elements in *Tex19.1^-/-^* spermatocytes causes a similar re-distribution of meiotic DSBs. Neither is it clear if loss of *Tex19.1* perturbs DSB formation at all meiotic recombination hotspots equally, or if any potential feedback mechanism stimulating DSBs on asynapsed regions of chromosomes influences DSB distribution. Thus, we cannot rule out the possibility that altered distribution of DSBs is contributing to chromosome asynapsis in *Tex19.1^-/-^* spermatocytes. However, DSB frequency in *Tex19.1^-/-^* spermatocytes is reduced to a level similar to that seen in hypomorphic *Spo11* mutants, which also have defects in chromosome synapsis [14,33]. Thus, reduced DSB frequency is likely the primary cause of chromosome asynapsis in *Tex19.1^-/-^* spermatocytes.

### Roles for *Tex19.1* and *Ubr2* in Meiotic DSB Formation

The data presented here shows that both *Tex19.1* and *Ubr2* are required to generate sufficient meiotic DSBs in spermatocytes to ensure robust chromosome synapsis. UBR2 was previously suggested not to have a role in the initiation of meiotic recombination as it did not localise to recombination foci and was not required for normal recruitment of RAD51 or RPA to recombination foci during leptotene [42]. Immunocytologically-detectable enrichment at recombination foci is probably not a requirement for UBR2 to directly or indirectly influence the initiation of meiotic recombination, however the effect of *Ubr2* on recombination foci and γH2AX during leptotene reported here does contradict statements in that previous report [42]. Differences between mouse strain background or *Ubr2* allele being studied may contribute to this, and the delay in synapsis initiation during zygotene (Supporting Fig S2) could also complicate meiotic prophase substaging during analysis of *Ubr2*^*-/-*^ spermatocytes leading to differences between studies. Furthermore, representative images and quantitative analysis of recombination foci in leptotene *Ubr2*^*-/-*^ spermatocytes are not shown in the previous report [42], making it difficult to directly compare data between studies. However reduced numbers of recombination foci in zygotene *Ubr2*^*-/-*^spermatocytes have been reported previously [42] and are consistent with the data presented here. Our data indicates that the reduction in zygotene recombination foci in *Ubr2*^*-/-*^ spermatocytes is a consequence of earlier defects in the generation of meiotic DSBs, and that reduced initiation of meiotic recombination is contributing to the *Ubr2*^*-/-*^ meiotic phenotype. This aspect of the *Ubr2*^*-/-*^phenotype may be caused by the absence of detectable TEX19.1 protein in *Ubr2*^*-/-*^ testes.

The phenotypic similarity between *Tex19.1^-/-^* mutants and *Ubr2*^*-/-*^ mutants, in combination with the physical interaction between TEX19.1 and UBR2 proteins [25], and the requirement for *Ubr2* for TEX19.1 protein stability [25], suggests that TEX19.1 and UBR2 are functioning in the same pathway to promote meiotic DSB formation. However, the molecular mechanism underlying the genetic requirement for *Tex19.1* and *Ubr2* in robust initiation of meiotic recombination is not clear. It is possible that the defects in the initiation of meiotic recombination in *Ubr2*^*-/-*^ spermatocytes described here reflects the absence of TEX19.1 protein in these cells and uncharacterised downstream functions of TEX19.1 in regulating meiotic recombination. Alternatively, It is possible that TEX19.1 is regulating the activity of UBR2, a RING-domain E3 ubiquitin ligase, and that TEX19.1’s effects on meiotic DSB formation reflect a role for at least one of UBR2’s substrates in meiotic recombination. Indeed, loss of *Tex19.1* might have effects on multiple UBR2 substrates and that could be responsible for the different aspects of the *Tex19.1^-/-^* phenotype in different developmental stages and tissues. UBR2 has been shown to ubiquitylate histone H2A and histone H2B [41], and is implicated in degrading the C-terminal fragment of the REC8 cohesin subunit generated by separase-dependent cleavage [43]. Therefore it is possible that loss of *Tex19.1* or *Ubr2* affects meiotic chromosome organisation or the meiotic chromatin substrate on which the SPO11 endonuclease is acting. It is also possible that UBR2 ubiquitylates SPO11, or one of its regulators, and that loss of *Tex19.1* or *Ubr2* affects the amount of SPO11 activity rather than its chromatin substrate in leptotene spermatocytes. These possibilities are not mutually exclusive and further work is required to elucidate how the *Tex19.1*-*Ubr2* pathway influences DSB formation. However, the data presented here demonstrates that *Tex19.1* and *Ubr2* are genetically required to ensure that sufficient meiotic DSBs are generated in male meiosis.

## Materials and Methods

### Mice

*Tex19.1^-/-^* animals on a C57BL/6 genetic background were bred and genotyped as described [23]. *Spo11*^*+/-*^ heterozygous mice [7] on a C57BL/6 genetic background [18] were inter-crossed with *Tex19.1^+/-^* mice. Animal experiments were carried out under UK Home Office Project Licence PPL 60/4424. Day of birth was designated P1, adult mice were typically analysed at between 6-14 weeks old. *Tex19.1^+/+^* and *Tex19.1^+/-^* animals have no difference in sperm counts [23] and data from these control animals from the same breeding colony were pooled as *Tex19.1^+/±^*. Epididymal sperm counts were determined as described [23]. *Ubr2^-/-^* mice were generated by CRISPR/Cas9 double nickase-mediated genome editing in zygotes [44]. Complementary oligonucleotides (Supporting Table S2) targeting exon 3 of *Ubr2* were annealed and cloned into plasmid pX335 [45], amplified by PCR, then in vitro transcribed using a T7 Quick High Yield RNA Synthesis kit (NEB) to generate paired guide RNAs. RNA encoding the Cas9 nickase mutant (50 ng/µl, Tebu-Bio), paired guide RNAs targeting exon 3 of UBR2 (each at 25 ng/µl), and 150 ng/µl single-stranded DNA oligonucleotide repair template (Supporting Table S2) were microinjected into the cytoplasm of C57BL/6 × CBA F2 zygotes. The repair template introduces an XbaI restriction site and mutates cysteine-121 within the UBR domain of *Ubr2* (Uniprot Q6WKZ8-1) to a premature stop codon. The zygotes were then cultured overnight in KSOM (Millipore) and transferred into the oviduct of pseudopregnant recipient females. Pups were genotyped and the mutant *Ubr2* allele back-crossed to C57BL/6.

### Immunostaining Meiotic Chromosome Spreads

Chromosome spreads were prepared as described by Peters et al. [46] for the *Spo11*^*-/-*^ and *Tex19.1^-/-^Spo11^-/-^* analyses, or by Costa et al. [47] for all other analyses. For immunostaining, slides were blocked and antibodies diluted in PBS containing 0.15% BSA, 0.1% Tween-20 and 5% goat serum as indicated in Supporting Table 1. The anti-MEI4, anti-SYCE2, and anti-RPA primary antibodies used were as reported [34,48,49]. Alexa Fluor-conjugated secondary antibodies (Invitrogen) were used at a 1:500 dilution, and 2 ng/μl 4,6-diamidino-2-phenylidole (DAPI) was used to fluorescently stain DNA. Slides were mounted in 90% glycerol, 10% PBS, 0.1% p-phenylenediamine. Three or four channel images were captured with iVision or IPLab software (BioVision Technologies) using an Axioplan II fluorescence microscope (Carl Zeiss) equipped with motorised colour filters. Immunostaining was performed on spreads from at least three experimental and three control animals. Statistical analysis was performed in R [50], means are reported ± standard error, and n is reported as total number of spreads analysed in each experiment.

Nuclei were staged by immunostaining for the axial/lateral element marker SYCP3 [51]. Nuclei with short fragments of axial element were classified as leptotene, those with complete axial filaments undergoing synapsis as zygotene, and those with complete autosomal synapsis as pachytene. Immunostaining for the central element component SYCE2 [47], or the transverse filament component SYCP1 [52], were included in some experiments to monitor synapsis.

Asynapsed pachytene *Tex19.1^-/-^* nuclei [23] were distinguished from zygotene due to incomplete sets of synapsed autosomes. Recombination foci in leptotene and zygotene nuclei were imaged by capturing z-stacks using a piezoelectrically-driven objective mount (Physik Instrumente) controlled with Volocity software (PerkinElmer). These images were deconvolved using Volocity, a 2D image generated in Fiji [53], and analysed in Adobe Photoshop CS6. DMC1, RAD51 and RPA foci were counted as recombination foci when they overlapped a chromosome axis. To measure leptotene γH2AX or H3K4me3 signal intensity, nuclear area was delimited using the DAPI signal, and signal intensity in that area quantified and corrected for background non-nuclear signal in 16 bit grayscale images using Fiji software. To assess the extent of synapsis in zygotene nuclei, the length of completely assembled synaptonemal complex was estimated relative to the length of axial/lateral element staining in that nucleus. For this and all immunocytological scoring, images were scored blind with respect to genotype by pooling control and knockout images, randomly assigning new filenames to each image, then decoding the filenames after scoring.

### Chromatin Immunoprecipitation (ChIP)

Decapsulated P16 testes were macerated with razor blades in ice-cold PBS, tissue fragments were removed by allowing to settle, and testicular cells pelleted at 860*g* for 5 minutes at 4°C. The cells were resuspended in PBS and cross-linking ChIP performed essentially as described [54]. 5µl rabbit anti-histone H3K4me3 antibody (Millipore) was coupled to 20µl Dynabeads-Protein A (Life Technologies) for each ChIP. DNA was purified using MinElute PCR Purification Kits (Qiagen), eluted in 20µl buffer EB, and diluted 1:10 for quantitative PCR (qPCR) using SYBR Select Master Mix (Applied Biosystems). ChIP and input samples from three biological replicates of *Tex19.1*^*+/±*^and *Tex19.1^-/-^* P16 testes were assayed in triplicate by qPCR using SYBR Green Master Mix (Roche) and a LightCycler 480 (Roche). ChIP enrichment was calculated relative to 10% input samples, and normalised to enrichment for the β-actin (*Actb*) transcriptional start site. Primers used for qPCR are listed in Supporting Table S2. Primers for *Prdm9*-dependent recombination hotspots were derived from DMC1 ChIP-seq data [5].

### Histology and In Situ Hybridisation

Histology of Bouin’s-fixed testes, and in situ hybridisation of MMERVK10C probes to Bouin’s-fixed testis sections were performed as described [23].

### qRT-PCR

RNA was isolated from macerated mouse testes using TRIzol (Invitrogen) and treated with Turbo DNAse (Ambion) to digest any genomic DNA contamination. 1µg DNAse-treated RNA was used to synthesise cDNA using Superscript III (Invitrogen). The cDNA was used as a template for qPCR using SYBR Select Master Mix (Applied Biosystems), and the relative quantity of RNA transcript calculated using the standard curve method as described by the supplier. The qPCR was performed on the LightCycler 480 (Roche), retrotransposon RNA levels were measured relative to β-actin, and normalised to control samples. Each biological replicate was assayed in triplicate, and alongside no reverse transcriptase and no template control reactions to confirm the absence of genomic DNA contamination.

### Western Blotting

P16 testes were homogenised in 2× Laemmli SDS sample buffer (Sigma) with a motorised pestle, boiled for 2-5 minutes and insoluble material pelleted in a microcentrifuge. Lysates were resolved by electrophoresis through pre-cast Bis-Tris polyacrylamide gels (Life Technologies) and Western blotted to PVDF membrane using the iBlot Dry Blotting System (Life Technologies). PBS containing 5% skimmed milk and 0.1% Tween was used to block the membrane and dilute antibodies. Primary antibodies for Western blotting were mouse anti-UBR2 (Abcam, 1:1000 dilution) and mouse anti-β-actin (Sigma, 1:5000 dilution). HRP-conjugated secondary antibodies (Cell Signaling Technology; Bio-Rad) were detected with SuperSignal West Pico Chemiluminescent Substrate (Thermo Scientific).

## Acknowledgements

We thank Bernard de Massy (IGH, Montpellier, France) and C. James Ingles (University of Toronto, Canada) for anti-MEI4 and anti-RPA antibodies respectively, James Turner (MRC NIMR, London, UK) and Bernard de Massy for *Spo11* mutant mice, Pradeepa Madapura (MRC HGU, Edinburgh, UK) for help with ChIP experiments, and Chao-Chun Hung and Judith Reichmann (both MRC HGU, Edinburgh, UK) for *Tex19.1* expression clones and reagents. We thank Wendy Bickmore and Javier Caceres (both MRC HGU, Edinburgh, UK) for advice and comments on the manuscript.

## References

1. HandelMA, SchimentiJC. Genetics of mammalian meiosis: regulation, dynamics and impact on fertility. Nat Rev Genet. 2010;11: 124–136. doi:10.1038/nrg2723

2. BaudatF, ImaiY, de MassyB. Meiotic recombination in mammals: localization and regulation. Nat Rev Genet. 2013;14: 794–806. doi: 10.1038/nrg3573

3. BordeV, de MassyB., Programmed induction of DNA double strand breaks during meiosis: setting up communication between DNA and the chromosome structure. Curr Opin Genet Dev. 2013;23: 147–155. doi: 10.1016/j.gde.2012.12.002

4. BaudatF, BuardJ, GreyC, Fledel-AlonA, OberC, PrzeworskiM, et al. PRDM9 is a major determinant of meiotic recombination hotspots in humans and mice. Science. 2010;327: 836–840. doi: 10.1126/science.1183439

5. BrickK, SmagulovaF, KhilP, Camerini-OteroRD, PetukhovaGV. Genetic recombination is directed away from functional genomic elements in mice. Nature. 2012;485: 642–645. doi: 10.1038/nature11089

6. RobertT, NoreA, BrunC, MaffreC, CrimiB, BourbonH-M, et al. The TopoVIB-Like protein family is required for meiotic DNA double-strand break formation. Science. 2016;351: 943–949. doi: 10.1126/science.aad5309

7. BaudatF, ManovaK, YuenJP, JasinM, KeeneyS., Chromosome synapsis defects and sexually dimorphic meiotic progression in mice lacking Spo11. Mol Cell. 2000;6: 989–998. doi: 10.1016/S1097-2765(00)00098-8

8. RomanienkoPJ, Camerini-OteroRD., The mouse Spo11 gene is required for meiotic chromosome synapsis. Mol Cell. 2000;6: 975–987. doi: 10.1016/S1097-2765(00)00097-6.

9. MahadevaiahSK, TurnerJM, BaudatF, RogakouEP, de BoerP, Blanco-RodríguezJ, et al. Recombinational DNA double-strand breaks in mice precede synapsis. Nat Genet. 2001;27: 271–276. doi: 10.1038/85830

10. NealeMJ, PanJ, KeeneyS., Endonucleolytic processing of covalent protein-linked DNA double-strand breaks. Nature. 2005;436: 1053–1057. doi: 10.1038/nature03872

11. BellaniMA, BoatengKA, McLeodD, Camerini-OteroRD. The expression profile of the major mouse SPO11 isoforms indicates that SPO11beta introduces double strand breaks and suggests that SPO11alpha has an additional role in prophase in both spermatocytes and oocytes. Mol Cell Biol. 2010;30: 4391–4403. doi: 10.1128/MCB.00002-10

12. KauppiL, BarchiM, BaudatF, RomanienkoPJ, KeeneyS, JasinM., Distinct properties of the XY pseudoautosomal region crucial for male meiosis. Science. 2011;331: 916–920. doi: 10.1126/science.1195774

13. LangeJ, PanJ, ColeF, ThelenMP, JasinM, KeeneyS. ATM controls meiotic double-strand-break formation. Nature. 2011;479: 237–240. doi: 10.1038/nature10508

14. KauppiL, BarchiM, LangeJ, BaudatF, JasinM, KeeneyS. Numerical constraints and feedback control of double-strand breaks in mouse meiosis. Genes Dev. 2013;27: 873–886. doi: 10.1101/gad.213652.113

15. CrichtonJH, DunicanDS, MacLennanM, MeehanRR, AdamsIR. Defending the genome from the enemy within: mechanisms of retrotransposon suppression in the mouse germline. Cell Mol Life Sci. 2014;71: 1581–1605. doi: 10.1007/s00018-013-1468-0

16. SoperSFC, van der HeijdenGW, HardimanTC, GoodheartM, MartinSL, de BoerP, et al. Mouse maelstrom, a component of nuage, is essential for spermatogenesis and transposon repression in meiosis. Dev Cell. 2008;15: 285–297. doi: 10.1016/j.devcel.2008.05.015

17. Bourc’hisD, BestorTH., Meiotic catastrophe and retrotransposon reactivation in male germ cells lacking Dnmt3L. Nature. 2004;431: 96–99. doi: 10.1038/nature02886

18. MahadevaiahSK, Bourc’hisD, de RooijDG, BestorTH, TurnerJMA, BurgoynePS. Extensive meiotic asynapsis in mice antagonises meiotic silencing of unsynapsed chromatin and consequently disrupts meiotic sex chromosome inactivation. J Cell Biol. 2008;182: 263–276. doi: 10.1083/jcb.200710195

19. ZamudioN, BarauJ, TeissandierA, WalterM, BorsosM, ServantN, et al. DNA methylation restrains transposons from adopting a chromatin signature permissive for meiotic recombination. Genes Dev. 2015;29: 1256–1270. doi: 10.1101/gad.257840.114

20. OllingerR, ReichmannJ, AdamsIR. Meiosis and retrotransposon silencing during germ cell development in mice. Differentiation. 2010;79: 147–158. doi: 10.1016/j.diff.2009.10.004

21. WangPJ, McCarreyJR, YangF, PageDC., An abundance of X-linked genes expressed in spermatogonia. Nat Genet. 2001;27: 422–426. doi: 10.1038/86927

22. KuntzS, KiefferE, BianchettiL, LamoureuxN, FuhrmannG, VivilleS. Tex19, a mammalian-specific protein with a restricted expression in pluripotent stem cells and germ line. Stem Cells. 2008;26: 734–744. doi: 10.1634/stemcells.2007-0772

23. ÖllingerR, ChildsAJ, BurgessHM, SpeedRM, LundegaardPR, ReynoldsN, et al. Deletion of the pluripotency-associated Tex19.1 gene causes activation of endogenous retroviruses and defective spermatogenesis in mice. PLoS Genet. 2008;4: e1000199. doi: 10.1371/journal.pgen.1000199

24. HackettJA, ReddingtonJP, NestorCE, DunicanDS, BrancoMR, ReichmannJ, et al. Promoter DNA methylation couples genome-defence mechanisms to epigenetic reprogramming in the mouse germline. Development. 2012;139: 3623–3632. doi: 10.1242/dev.081661

25. YangF, ChengY, AnJY, KwonYT, EckardtS, LeuNA, et al. The ubiquitin ligase Ubr2, a recognition E3 component of the N-end rule pathway, stabilizes Tex19.1 during spermatogenesis. PLoS ONE. 2010;5: e14017. doi: 10.1371/journal.pone.0014017

26. YoshidaK, KondohG, MatsudaY, HabuT, NishimuneY, MoritaT., The mouse RecA-like gene Dmc1 is required for homologous chromosome synapsis during meiosis. Mol Cell. 1998;1: 707–718. doi: 10.1016/S1097-2765(00)80070-2

27. de VriesFAT, de BoerE, van den BoschM, BaarendsWM, OomsM, YuanL, et al. Mouse Sycp1 functions in synaptonemal complex assembly, meiotic recombination, and XY body formation. Genes Dev. 2005;19: 1376–1389. doi: 10.1101/gad.329705

28. DanielK, LangeJ, HachedK, FuJ, AnastassiadisK, RoigI, et al. Meiotic homologue alignment and its quality surveillance are controlled by mouse HORMAD1. Nat Cell Biol. 2011;13: 599–610. doi: 10.1038/ncb2213

29. MoensPB, KolasNK, TarsounasM, MarconE, CohenPE, SpyropoulosB., The time course and chromosomal localization of recombination-related proteins at meiosis in the mouse are compatible with models that can resolve the early DNA-DNA interactions without reciprocal recombination. J Cell Sci. 2002;115: 1611–1622.

30. KidaneD, JonasonAS, GortonTS, MihaylovI, PanJ, KeeneyS, et al. DNA polymerase beta is critical for mouse meiotic synapsis. EMBO J. 2010;29: 410–423. doi: 10.1038/emboj.2009.357

31. ChicheporticheA, Bernardino-SgherriJ, de MassyB, DutrillauxB., Characterization of Spo11-dependent and independent phospho-H2AX foci during meiotic prophase I in the male mouse. J Cell Sci. 2007;120: 1733–1742. doi: 10.1242/jcs.004945

32. CarofiglioF, InagakiA, de VriesS, WassenaarE, SchoenmakersS, VermeulenC, et al. SPO11-independent DNA repair foci and their role in meiotic silencing. PLoS Genet. 2013;9: e1003538. doi: 10.1371/journal.pgen.1003538

33. ColeF, KauppiL, LangeJ, RoigI, WangR, KeeneyS, et al. Homeostatic control of recombination is implemented progressively in mouse meiosis. Nat Cell Biol. 2012;14: 424–430. doi: 10.1038/ncb2451

34. 37. KumarR, BourbonH-M, de MassyB., Functional conservation of Mei4 for meiotic DNA double-strand break formation from yeasts to mice. Genes Dev. 2010;24: 1266–1280. doi: 10.1101/gad.571710

35. HayashiK, YoshidaK, MatsuiY., A histone H3 methyltransferase controls epigenetic events required for meiotic prophase. Nature. 2005;438: 374–378. doi: 10.1038/nature04112

36. SunF, FujiwaraY, ReinholdtLG, HuJ, SaxlRL, BakerCL, et al. Nuclear localization of PRDM9 and its role in meiotic chromatin modifications and homologous synapsis. Chromosoma. 2015; doi: 10.1007/s00412-015-0511-3

37. CrichtonJH, PlayfootCJ, AdamsIR. The role of chromatin modifications in progression through mouse meiotic prophase. J Genet Genomics. 2014;41: 97–106. doi: 10.1016/j.jgg.2014.01.003

38. ReichmannJ, CrichtonJH, MadejMJ, TaggartM, GautierP, Garcia-PerezJL, et al. Microarray analysis of LTR retrotransposon silencing identifies Hdac1 as a regulator of retrotransposon expression in mouse embryonic stem cells. PLoS Comput Biol. 2012;8: e1002486. doi: 10.1371/journal.pcbi.1002486

39. ReichmannJ, ReddingtonJP, BestD, ReadD, ÖllingerR, MeehanRR, et al. The genome-defence gene Tex19.1 suppresses LINE-1 retrotransposons in the placenta and prevents intra-uterine growth retardation in mice. Hum Mol Genet. 2013;22: 1791–1806. doi: 10.1093/hmg/ddt029

40. KwonYT, XiaZ, AnJY, TasakiT, DavydovIV, SeoJW, et al. Female lethality and apoptosis of spermatocytes in mice lacking the UBR2 ubiquitin ligase of the N-end rule pathway. Mol Cell Biol. 2003;23: 8255–8271.

41. AnJY, KimE-A, JiangY, ZakrzewskaA, KimDE, LeeMJ, et al. UBR2 mediates transcriptional silencing during spermatogenesis via histone ubiquitination. Proc Natl Acad Sci USA. 2010;107: 1912–1917. doi: 10.1073/pnas.0910267107

42. AnJY, KimE, ZakrzewskaA, YooYD, JangJM, HanDH, et al. UBR2 of the N-end rule pathway is required for chromosome stability via histone ubiquitylation in spermatocytes and somatic cells. PLoS ONE. 2012;7: e37414. doi: 10.1371/journal.pone.0037414

43. LiuY-J, LiuC, ChangZ, WadasB, BrowerCS, SongZ-H, et al. Degradation of the Separase-cleaved Rec8, a Meiotic Cohesin Subunit, by the N-end Rule Pathway. J Biol Chem. 2016;291: 7426–7438. doi: 10.1074/jbc.M116.714964

44. RanFA, HsuPD, LinC-Y, GootenbergJS, KonermannS, TrevinoAE, et al. Double nicking by RNA-guided CRISPR Cas9 for enhanced genome editing specificity. Cell. 2013;154: 1380–1389. doi: 10.1016/j.cell.2013.08.021

45. CongL, RanFA, CoxD, LinS, BarrettoR, HabibN, et al. Multiplex genome engineering using CRISPR/Cas systems. Science. 2013;339: 819–823. doi: 10.1126/science.1231143

46. PetersAH, PlugAW, van VugtMJ, de BoerP. A drying-down technique for the spreading of mammalian meiocytes from the male and female germline. Chromosome Res. 1997;5: 66–68. doi: 10.1023/A:1018445520117

47. CostaY, SpeedR, ÖllingerR, AlsheimerM, SempleCA, GautierP, et al. Two novel proteins recruited by synaptonemal complex protein 1 (SYCP1) are at the centre of meiosis. J Cell Sci. 2005;118: 2755–2762. doi: 10.1242/jcs.02402

48. HeZ, HenricksenLA, WoldMS, InglesCJ. RPA involvement in the damage-recognition and incision steps of nucleotide excision repair. Nature. 1995;374: 566–569. doi: 10.1038/374566a0

49. Bolcun-FilasE, HallE, SpeedR, TaggartM, GreyC, de MassyB, et al. Mutation of the mouse Syce1 gene disrupts synapsis and suggests a link between synaptonemal complex structural components and DNA repair. PLoS Genet. 2009;5: e1000393. doi: 10.1371/journal.pgen.1000393

50. R Core Team. R: A language and environment for statistical computing. R Foundation for Statistical Computing, Vienna, Austria. [Internet]. 2014. Available: http://www.r-project.org/

51. LammersJH, OffenbergHH, van AalderenM, VinkAC, DietrichAJ, HeytingC., The gene encoding a major component of the lateral elements of synaptonemal complexes of the rat is related to X-linked lymphocyte-regulated genes. Mol Cell Biol. 1994;14: 1137–1146.

52. MeuwissenRL, OffenbergHH, DietrichAJ, RiesewijkA, van IerselM, HeytingC. A coiled-coil related protein specific for synapsed regions of meiotic prophase chromosomes. EMBO J. 1992;11: 5091–5100.

53. SchindelinJ, Arganda-CarrerasI, FriseE, KaynigV, LongairM, PietzschT, et al. Fiji: an open-source platform for biological-image analysis. Nat Methods. 2012;9: 676–682. doi: 10.1038/nmeth.2019

54. MortazaviA, Leeper ThompsonEC, GarciaST, MyersRM, WoldB., Comparative genomics modeling of the NRSF/REST repressor network: from single conserved sites to genome-wide repertoire. Genome Res. 2006;16: 1208–1221. doi: 10.1101/gr.4997306

